# A *ROR2* coding variant is associated with craniofacial variation in domestic pigeons

**DOI:** 10.1101/2021.03.15.435542

**Authors:** Elena F. Boer, Hannah F. Van Hollebeke, Carson Holt, Mark Yandell, Michael D. Shapiro

## Abstract

Vertebrate craniofacial morphogenesis is a highly orchestrated process that is directed by evolutionarily conserved developmental pathways ^1,2^. Within species, canalized developmental programs typically produce only modest morphological variation. However, as a result of millennia of artificial selection, the domestic pigeon (*Columba livia*) displays radical variation in craniofacial morphology within a single species. One of the most striking cases of pigeon craniofacial variation is the short beak phenotype, which has been selected in numerous breeds. Classical genetic experiments suggest that pigeon beak length is regulated by a small number of genetic factors, one of which is sex-linked (*Ku2* locus) ^3–5^. However, the molecular genetic underpinnings of pigeon craniofacial variation remain unknown. To determine the genetic basis of the short beak phenotype, we used geometric morphometrics and quantitative trait loci (QTL) mapping on an F_2_ intercross between a short-beaked Old German Owl (OGO) and a medium-beaked Racing Homer (RH). We identified a single locus on the Z-chromosome that explains a majority of the variation in beak morphology in the RH x OGO F_2_ population. In complementary comparative genomic analyses, we found that the same locus is also strongly differentiated between breeds with short and medium beaks. Within the differentiated *Ku2* locus, we identified an amino acid substitution in the non-canonical Wnt receptor ROR2 as a putative regulator of pigeon beak length. The non-canonical Wnt (planar cell polarity) pathway serves critical roles in vertebrate neural crest cell migration and craniofacial morphogenesis ^6,7^. In humans, homozygous *ROR2* mutations cause autosomal recessive Robinow syndrome, a rare congenital disorder characterized by skeletal abnormalities, including a widened and shortened facial skeleton ^8,9^. Our results illustrate how the extraordinary craniofacial variation among pigeons can reveal genetic regulators of vertebrate craniofacial diversity.

## Results and Discussion

The avian beak shows remarkable diversity among species. Variation in beak morphology within groups like Darwin’s finches and Hawaiian honeycreepers illustrates the diversifying potential of natural selection on beak skeletal structures and functions ^10,11^. Although the underlying genetic basis of the extraordinary variation among birds remains relatively poorly understood, several genes associated with overall beak size or linear dimensions of beak shape are known in a modest number of species (e.g., *COL4A5* in Great tits ^12^; *IGF1* in Black-bellied seedcrackers ^13^; *BMP4, CALM1, ALX1*, and *HMGA2* in Darwin’s finches ^14–18^). Unlike wild birds, the domestic pigeon beak is unconstrained by natural selection; the astounding level of morphological variation within this species is instead the product of intensive artificial selection. Some of the major axes of variation in craniofacial shape that distinguish distantly related avian species are recapitulated among breeds of domestic pigeon, despite different mechanisms of selection between wild and captive populations ^19,20^. Therefore, pigeons provide a unique opportunity to uncover genetic variants associated with the types of beak variation that exist throughout the radiation of birds.

### Pigeon beak length covaries with body size and braincase shape in an experimental cross

To determine the genetic architecture of beak length in pigeons, we established an F_2_ intercross between a male Racing Homer (RH) and a female Old German Owl (OGO). The RH, which “has been bred for one purpose – speed – almost to the exclusion of all other factors and traits” ^21^, has a medium-length beak that resembles the ancestral condition in rock pigeons (Figure 1A,B). In contrast, the OGO beak “is one of the distinctive characteristics” of the breed and is “short in appearance, which is partly caused by the broad width of the beak in relation to its length” ^22^ (Figure 1C,D).

**Figure 1.**
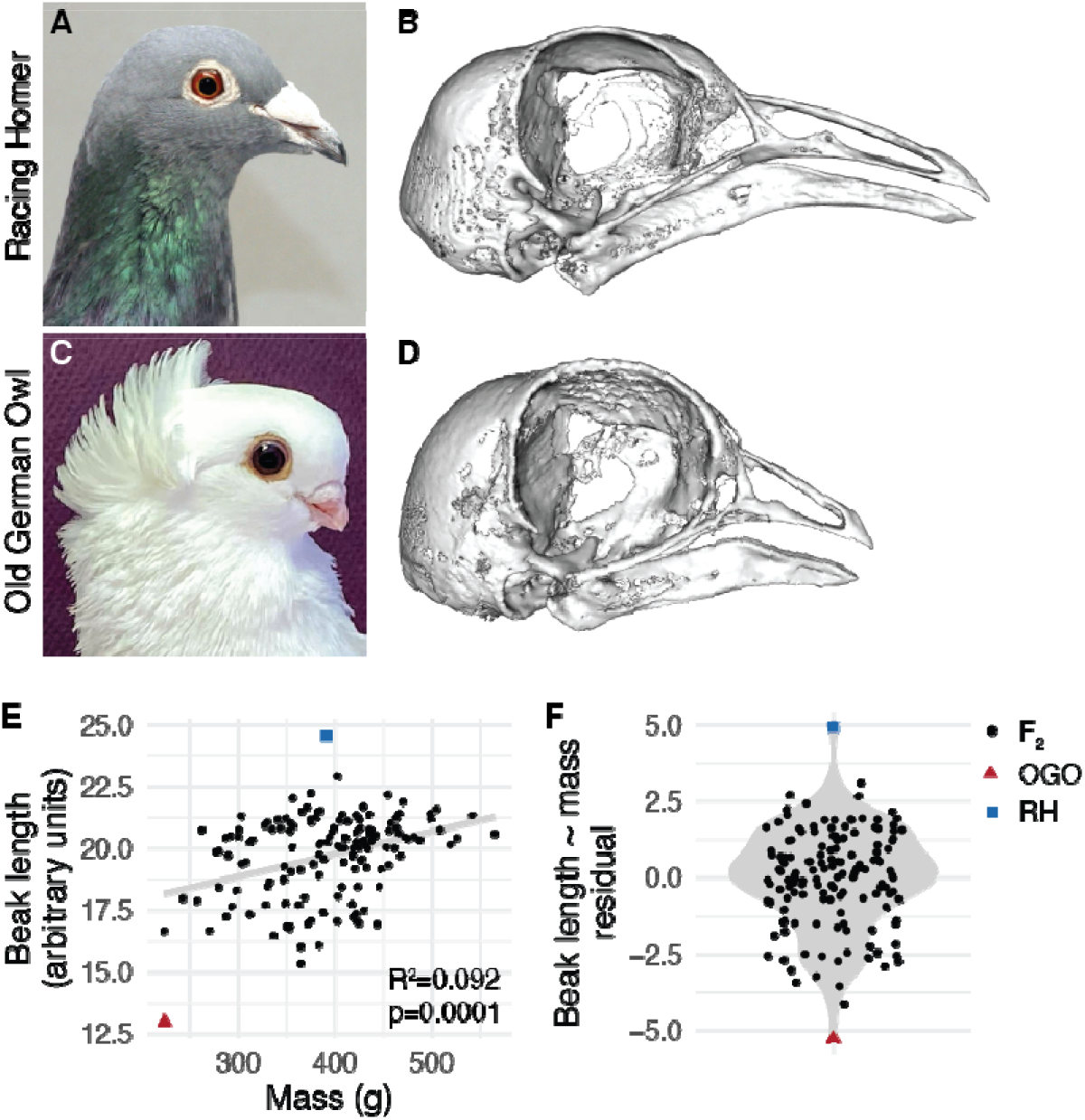
Beak length variation in a pigeon F_2_ intercross. (A) Representative image of the medium beak Racing Homer (RH) breed. (B) 3D surface model of the craniofacial skeleton of the male RH founder. (C) Representative image of the short beak Old German Owl (OGO) breed. (D) Surface model of the craniofacial skeleton of the female OGO founder. (E) Raw beak length (measured in arbitrary units) vs. total body mass (measured in grams) in the RH x OGO cross. Gray line indicates linear model from beak length ~ mass regression. (F) Distribution of residuals from beak length ~ mass regression. For (E-F), black dots denote RH x OGO F_2_ individuals, red triangle is OGO founder, blue square is RH founder. Image credits: Sydney Stringham (A), Brian McCormick (B).

We scanned the RH x OGO cross founders and 145 F_2_ individuals using micro-CT, generated 3D surface models of the craniofacial skeleton, and applied a set of 49 landmarks to the beak and braincase (Supplemental Figure 1, Supplemental Tables 1-2). By calculating the linear distance between the base and tip of the beak and using mass as a proxy for body size ^23^, we found a significant positive association between beak length and body size in the F_2_ population (R^2^=0.092, p=0.0001, Figure 1E). We removed the effects of body size variation by fitting a beak length ~ body size linear regression model and found that beak length residuals remained highly variable in the F_2_ population, demonstrating that beak length varies independently of body size (Figure 1F).

We also measured three-dimensional (3D) variation in beak and braincase shape through geometric morphometric analysis. In the RH x OGO F_2_ population, cranium centroid size is negatively associated with body size (R^2^=0.03, p=0.02, Supplemental Figure 2A). This result contrasts broad patterns of cranium ~ body size allometry observed across diverse pigeon breeds and wild birds ^20^, and is likely driven by the exceptionally large cranium and small body size selected for in the OGO breed. In the F_2_ population, cranium centroid size is also negatively associated with curvature from the tip of the beak to the back of the braincase and, to a lesser extent, beak length (R^2^=0.073, p<0.001, Supplemental Figure 2B). Taken together, the linear and 3D shape analyses reveal subtle but significant relationships between body and cranium size and craniofacial shape in the RH x OGO cross. By shuffling genetic programs for two generations in an experimental cross, we find that body size, cranium size, and beak shape are modular and separable.

Although allometry is an important correlate of shape ^23^, we focused further analyses on non-allometric craniofacial shape variation. Principal components analysis (PCA) of geometric morphometric shape variables demonstrate that, in the RH x OGO F_2_ population, the first two principal components (PCs) together account for 50% of craniofacial shape variation (Supplemental Figure 3A). The principal axis of shape variation (PC1, 39.1% of shape variation) describes compound variation in beak length and braincase volume (Figure 2A, Supplemental Movie 1) and is strongly correlated with linear measurements of beak length (R^2^=0.64, p<2.2e-16, Supplemental Figure 3B). PC2 (10.9% of shape variation) is defined almost exclusively by changes in braincase shape (Supplemental Figure 3C, Supplemental Movie 2).

**Figure 2.**
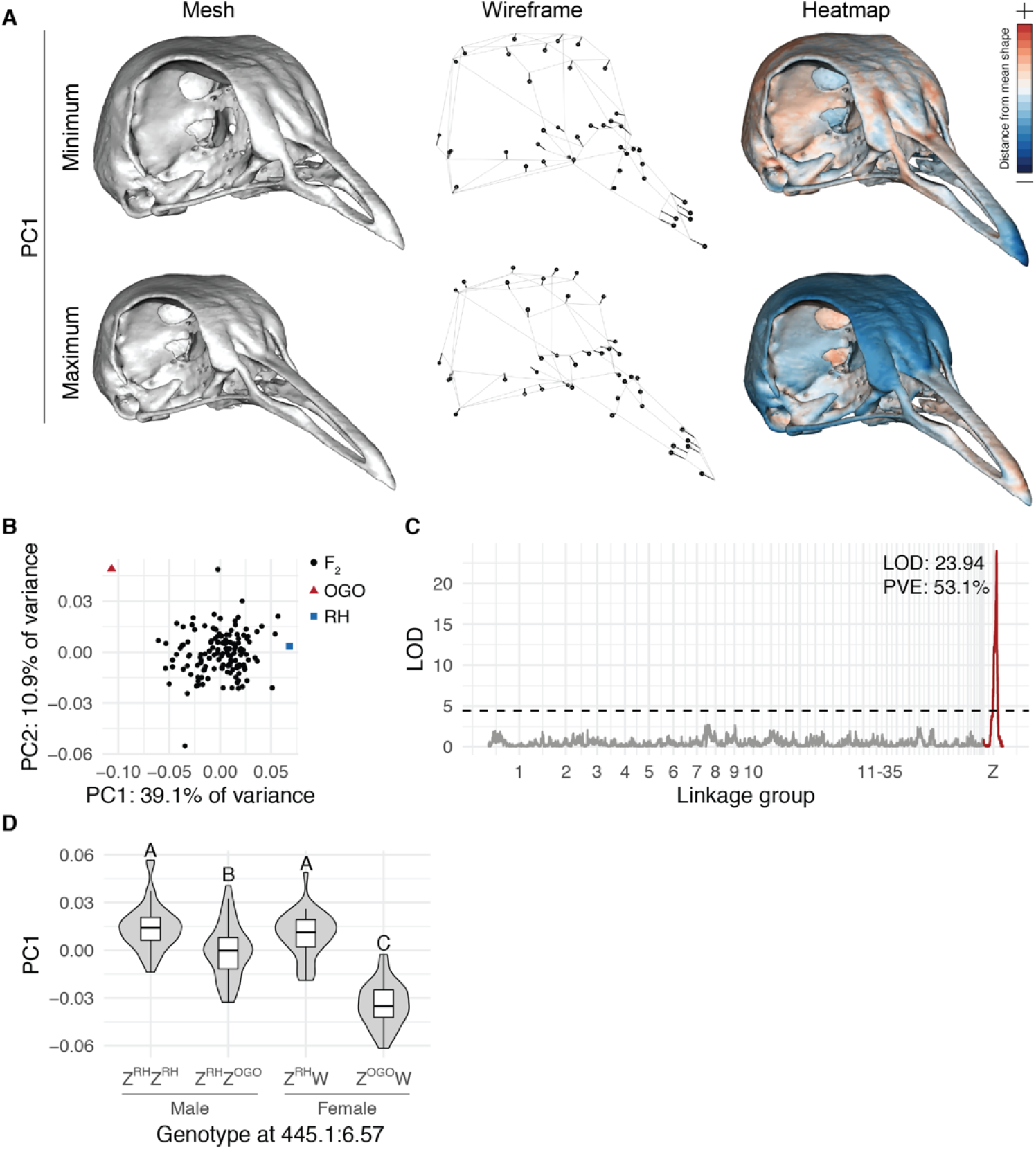
A major-effect QTL on the Z-chromosome is associated with principal component 1 (PC1) in the RH x OGO F2 population. (A) Visualizations of geometric morphometric PC1 minimum and maximum shapes three ways: warped 3D surface meshes (left), wireframes showing displacement of landmarks from mean shape (center), and heatmaps displaying regional shape variation (right). For warped meshes and wireframes, shape changes are magnified 1.5x to aid visualization. (B) PCA plot of PC1 vs. PC2. (C) Genome-wide QTL scan for PC1 reveals significant QTL on Z linkage group. (D) PC1 effect plot estimated from QTL peak marker. Letters denote significance groups; RH = allele from RH founder, OGO = allele from OGO founder.

Similar to previous findings in domestic pigeons and wild birds ^20,24^, 3D beak and braincase shape are strongly integrated in the RH x OGO F_2_ population (r-PLS=0.923, p<0.001, Supplemental Figure 2B). Along the PC1 axis, all F_2_ individuals are confined to a morphospace defined by the cross founders, but cluster closer to the RH than the OGO (Figure 2B). This result is reminiscent of our analysis of pigeon beak curvature, in which F_2_ individuals derived from a straight-beaked Pomeranian Pouter and curved-beaked Scandaroon more closely resembled the Pomeranian Pouter and never achieved the extreme craniofacial curvature of the Scandaroon ^24^. Therefore, two different genetic crosses using four different pigeon breeds suggest that the most exaggerated versions of craniofacial traits require coordination of multiple genetic factors.

### Identification of a major-effect beak length QTL on the Z chromosome

Next, we used the PC1 scores, which primarily describe variation in beak length, to perform genome-wide QTL scans. A single major-effect QTL on the Z-chromosome linkage group was strongly associated with PC1 and explained more than half of phenotypic variance in the RH x OGO cross (log likelihood ratio (LOD)=23.72, percent variance explained (PVE)=53.2%, Figure 2C). Nearly identical results were obtained when beak length residuals were used for QTL mapping (Supplemental Figure 4). The identification of a major-effect QTL on the Z-chromosome is consistent with results from classical genetic studies that pointed to a sex-linked regulator of pigeon beak length ^3–5^.

We next used the peak marker to estimate QTL effects. Within the F_2_ population, male Z^RH^/Z^RH^ homozygotes and female Z^RH^/W hemizygotes had the highest PC1 scores (longest beaks) and were not statistically different from one another (Figure 2D). Male Z^RH^/Z^OGO^ heterozygotes had intermediate PC1 scores (Figure 2D), suggestive of an incompletely dominant pattern of inheritance. In contrast, female Z^OGO^/W hemizygotes had dramatically lower PC1 scores (shorter beaks) than all other F_2_ individuals carrying the RH allele (Figure 2D). Although the structure of our experimental cross did not generate homozygous Z^OGO^/Z^OGO^ males in the F_2_ generation, previous classical genetic studies have demonstrated that, in short-beaked pigeon breeds, body size is sex-associated but beak length is not ^5,29^. Therefore, we predict that the beaks of Z^OGO^/Z^OGO^ males would be indistinguishable from hemizygous Z^OGO^/W females, although we cannot rule out the possibility that an additional copy of the OGO allele could result in even shorter beaks in Z^OGO^/Z^OGO^ males.

In summary, our results support the model that pigeon beak length is a polygenic trait controlled largely by one sex-linked factor. Additional minor-effect QTL are likely modifying beak length in the RH x OGO cross, some of which may be detectable in a larger F_2_ population or an F_3_ generation that includes Z^OGO^/Z^OGO^ males.

### A ROR2 coding variant is associated with beak length across diverse domestic pigeon breeds

The beak length QTL represents a relatively large (3.6-Mb) genomic region that includes several genes expressed during pigeon craniofacial development (Supplemental Figure 5, Supplemental Table 3), thus limiting our ability to pinpoint the specific gene(s) and mutation(s) that regulate beak length in the RH x OGO cross. In addition, because the mapping population was derived from just two birds that represent a fraction of the morphological diversity across pigeon breeds, we have no way of knowing if the beak length QTL we identified is relevant beyond the RH x OGO cross. Short beaks are characteristic of numerous related pigeon breeds that belong to the Owl family, but have also been selected for in a variety of unrelated non-Owl breeds ^26–28^. Pigeon breeders might have repeatedly selected the same standing variant in different breeds, independent variants of the same gene in different breeds, or different genes altogether in different breeds.

To distinguish between independent and shared genetic origins of short beaks, we scanned for genomic variants associated with beak length across diverse pigeon breeds by comparing resequenced genomes of 56 short-beaked individuals from 31 (7 Owl and 24 non-Owl) breeds to 121 genomes from 58 medium- or long-beaked breeds and feral pigeons (Figure 3A-B). We then searched for genomic regions that were differentiated between these groups using two related differentiation statistics (wcF_ST_ ^29^ and pF_ST_ ^30^). A ~293-kb segment on the Z-chromosome scaffold ScoHet5_445.1 stood out as significantly differentiated between the short- and medium/long-beaked groups (top 0.1% by wcF_ST_; Figure 3C-D, Supplemental Figure 6) and was located within the genomic interval identified in our QTL scan. In the peak differentiated region, short-beaked pigeons displayed elevated levels of haplotype homozygosity relative to medium/long-beaked individuals, providing further support for widespread positive selection on this locus in short-beaked breeds (Figure 3E). Thus, the short beak allele identified in our QTL mapping experiments is not specific to either the OGO cross founder or the Owl family. Instead, the short beak allele on the Z-chromosome likely arose once and was repeatedly selected in different breeds.

**Figure 3.**
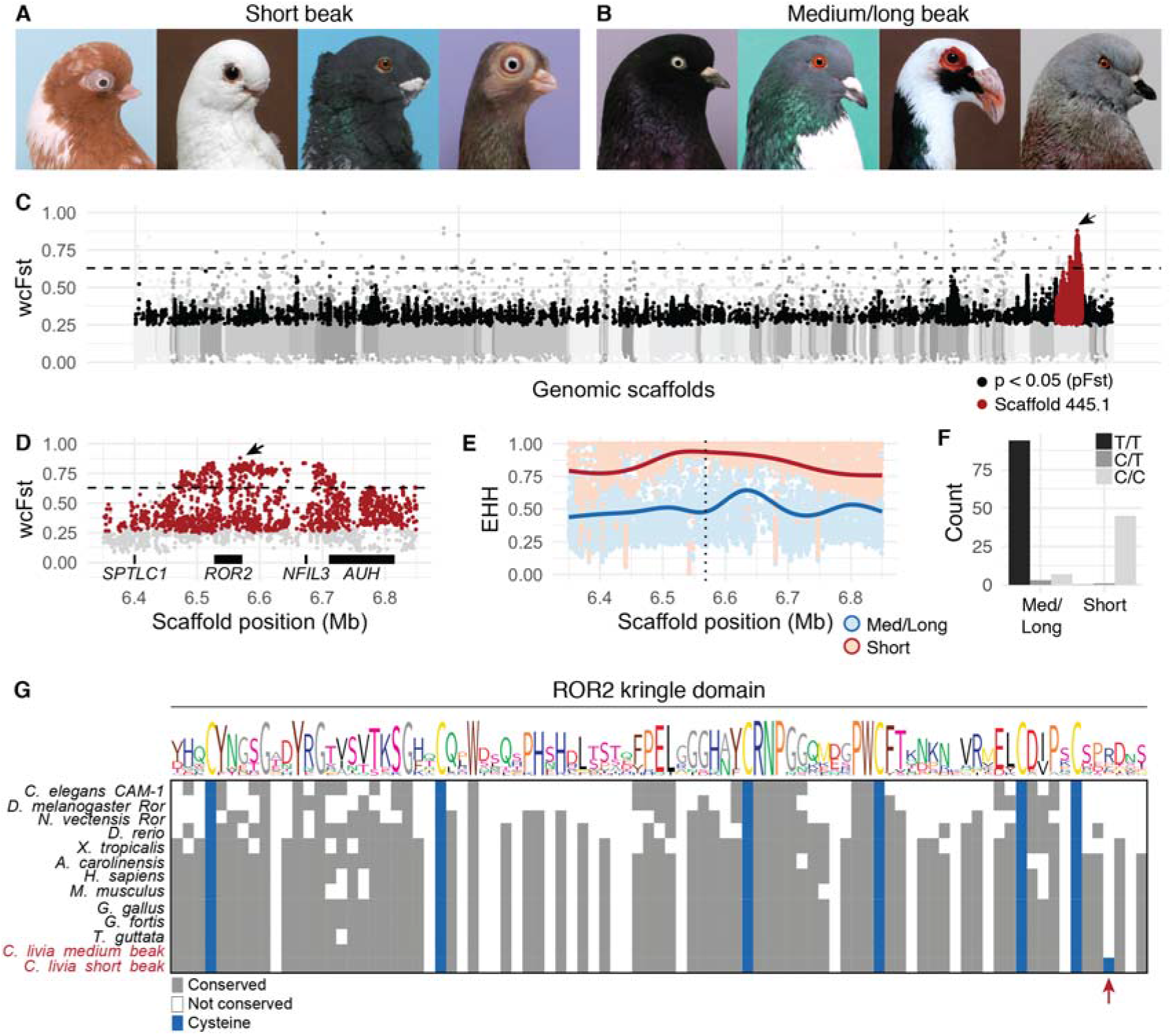
Comparison of short beak and medium/long beak pigeon genomes reveals *ROR2* coding variant. (A-B) Representative images of individuals representing short beak (A) and medium or long beak (B) pigeon breeds. (A) Short beak pigeons, from left to right: English Short Face Tumbler, African Owl, Oriental Frill, Budapest Tumbler. (B) Medium/long beak pigeons, from left to right: West of England, Cauchois, Scandaroon, Show King. (C) Genome-wide scan for allele frequency differentiation between short beak (n=56) and medium/long beak (n=121) pigeons. (D) Region of peak F_ST_ on ScoHet5_445.1; black horizontal bars represent four genes in the region. For (C-D), genomic scaffolds are colored in gray and ordered by genetic position in RH x OGO linkage map; black dots indicate SNPs that are significantly differentiated by pF_ST_ (Bonferroni-corrected p-value < 0.05); red dots are significant SNPs located on scaffold ScoHet5_445.1; dashed horizontal line represents threshold for genome-wide top 0.1% of differentiated SNPs by wcF_ST_; arrow points to ScoHet5_445.1:6568443, the most differentiated SNP (F_ST_=0.88) genome-wide. (E) Extended haplotype homozygosity in F_ST_ peak region; dotted vertical line indicates position of ScoHet5_445.1:6568443; smoothed lines represent local regression fitting ^54^. (F) Histogram of genotypes at ScoHet5_445.1:6568443 in short beak and medium/long beak groups. (G) Amino acid alignment of kringle domain from vertebrate ROR2 and invertebrate Ror homologs. The *Ku2* allele causes an arginine-to-cysteine substitution in short beak pigeon breeds.

The single most significantly-differentiated SNP genome wide (wcF_ST_=0.88, pF_ST_=0) is located at scaffold position ScoHet5_445.1:6568443. The non-reference allele causes a missense substitution in the seventh exon of *ROR2* (*ROR2^C1087T^*, hereafter the *Ku2* allele ^5^) in short-beaked pigeons. *ROR2* encodes a noncanonical Wnt receptor with well-established roles in cell polarity and motility in multiple embryonic tissues, including the neural crest ^31^. This gene is required for normal craniofacial development: in humans, homozygous mutations in *ROR2* cause autosomal recessive Robinow syndrome, a severe skeletal dysplasia characterized by extensive abnormalities, including a prominent forehead (frontal bossing), wideset eyes (hypertelorism), and a broad, short nose ^8,9^. In mice, *Ror2* knockout or knock-in of Robinow-associated mutations disrupts endochondral bone development and causes profound skeletal abnormalities, including craniofacial outgrowth defects ^32–34^. Likewise, the OGO and morphologically similar pigeon breeds have reduced craniofacial outgrowths in the form of short beaks.

Within the short-beaked group, 98% of pigeons (45/46) were homozygous or hemizygous for the *Ku2* allele; only the Chinese Nasal Tuft, a breed that can have a short- or medium-length beak ^35^, was heterogyzous. In contrast, 93% (97/104) of medium-beaked birds were homozygous, hemizygous, or heterozygous for the ancestral allele (Figure 3F). A genome-wide scan for putatively damaging coding variants predicted that the *Ku2* allele is both highly deleterious and associated with short beaks (VAAST ^36^ top-ranked feature, score=64.37, p=4e-8). At the amino acid level, the *Ku2* allele causes an arginine-to-cysteine transition in the ROR2 extracellular kringle fold, a cysteine-rich, disulfide-bonded domain that is unlikely to tolerate mutations due to its small size and complex folding ^37^. The precise number and spacing of cysteine residues in the kringle domain are deeply conserved in vertebrate ROR2 and invertebrate Ror homologs (Figure 3G), suggesting that the ectopic cysteine residue introduced by the *Ku2* mutation may have a substantial impact on disulfide bond formation and kringle domain folding. Although the precise function of the ROR2 kringle domain remains unclear, it is thought to mediate protein-protein interactions and may modulate the affinity of the adjacent Frizzled-like ligand-binding domain for WNT5A ^38,39^.

Like the pigeon *Ku2* allele, the majority of known Robinow-associated missense mutations in human patients are clustered in the kringle and Frizzled-like extracellular domains. All of the characterized disease variants cause increased ROR2 protein retention in the endoplasmic reticulum, suggesting that the extracellular domain must be properly folded before transport to the plasma membrane ^37,40^. Based on available evidence, we hypothesize that the *Ku2* allele disrupts ROR2 protein folding in short-beaked pigeons, resulting in craniofacial outgrowth anomalies similar to Robinow syndrome in humans.

### ROR2 and WNT5A are expressed during pigeon craniofacial morphogenesis

In chicken and mouse embryos, *ROR2* expression is widespread with regions of strong expression in the facial prominences, dorsal root ganglia, and limb buds ^31,41,42^. Using RNA-seq, we found that both *ROR2* and *WNT5A* are also strongly expressed in short- and medium-beaked pigeon facial primordia (n=5 each), with higher expression in the frontonasal and maxillary prominences (upper beak) relative to the mandibular prominence (lower beak; Figure 4A-B). Neither *ROR2* nor *WNT5A* is differentially expressed between short- and medium-beaked embryos at the pigeon equivalent of chicken stage HH29 (Ref. 43, Figure 4A-B), an early embryonic stage at which distinct craniofacial morphologies are evident among avian species ^44^.

**Figure 4.**
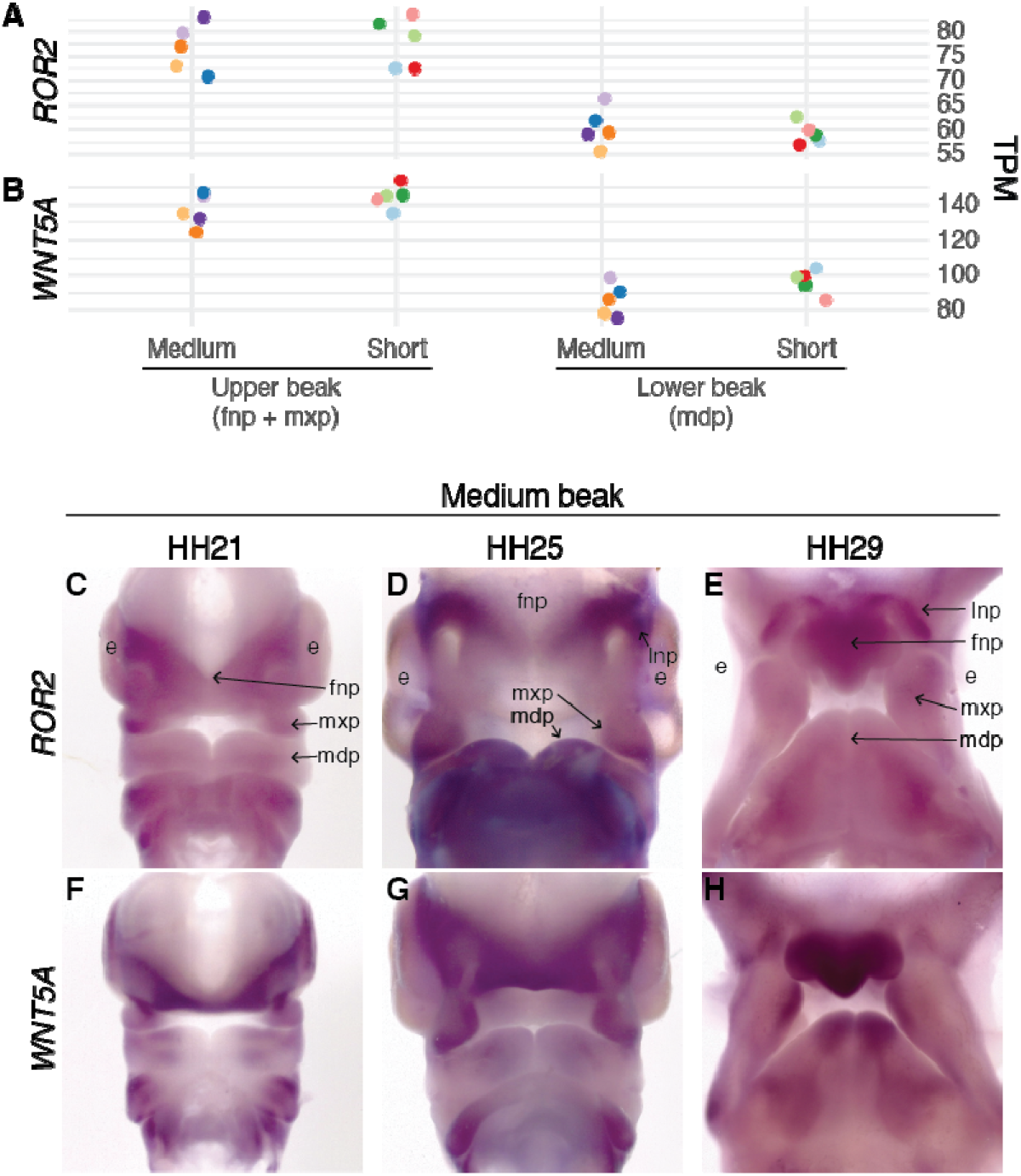
*ROR2* and *WNT5A* are expressed in pigeon facial primordia. (A-B) *ROR2* (A) and *WNT5A* (B) mRNA expression in facial primordia that will form the upper and lower beak from HH29 short and medium beak pigeon embryos. Each individual embryo is displayed in a different color. (C-H) Whole-mount *in situ* hybridization for *ROR2* (C-E) and *WNT5A* (F-H) in medium beak pigeon embryos at HH21, HH25, and HH29. *ROR2* is broadly expressed in facial primordia at all stages. *WNT5A* is strongly expressed in the FNP and at the lateral edges of the MXP and LNP, with increased expression at the edge of the MDP at HH29. Letters indicate embryonic tissues/structures: e=eye, fnp=frontonasal prominence, l=lateral nasal prominence, mdp=mandibular prominence, mxp=maxillary prominence.

Spatial expression of *ROR2* is broad during early pigeon facial development (HH21-29), with higher levels in the FNP and lateral nasal prominences (LNP) at HH25 and HH29 (Figure 4C-E). *WNT5A* expression domains overlap with *ROR2*, but are more spatially restricted to the regions of the facial primordia that will grow out to form the beak (Figure 4F-H), similar to mouse and chicken ^45,46^. Thus, *ROR2* and *WNT5A* are expressed together in pigeons in a spatial and temporal manner that is consistent with their role as regulators of craniofacial morphogenesis. The lack of differential *ROR2* expression in short- and medium-beaked pigeon embryos implicates the *Ku2* coding mutation, rather than differences in the regulation of expression, in the development of the short beak phenotype.

### Pigeons model vertebrate evolution and disease

Several developmental pathways have been implicated in the evolution of beak diversity in other birds, including Darwin’s finches, Great tits, and Black-bellied seedcrackers ^12–16^, Although *ROR2* has a well-established role in mammalian craniofacial development, to our knowledge, this is the first example of a member of the noncanonical Wnt signaling pathway regulating craniofacial development and diversity in birds. This finding contrasts with other examples of recurrent evolution of derived traits via changes in the same genes in pigeons and other species, including head crests (*EPHB2*, also in ringneck doves ^27,47^), feathered feet (*PITX1* and *TBX5*, also in chickens ^30,48^), and plumage color patterning (*NDP*, also in crows ^49–51^). Considering the deep evolutionary conservation of developmental pathways that regulate craniofacial morphogenesis, our findings raise the possibility that noncanonical Wnt signaling is modulated in other cases of avian craniofacial variation. We did not identify noncanonical Wnt pathway genes in our recent study of the genetic basis of pigeon beak elaboration ^24^, suggesting that distinct genetic programs underlie reduction and exaggeration of the same tissues and structures.

The identification of *ROR2* as a putative regulator of beak length adds to a growing list of genes that underlie morphological variation in the domestic pigeon and are associated with human diseases, including congenital defects and cancer ^30,50,52,53^. In addition, prior work in pigeons has predicted the molecular basis of diverse morphological traits in other wild and domestic species ^27,30,47,49–51^. Thus, the pigeon is an exceptional model to interrogate the genetic underpinnings of vertebrate evolution, development, and disease.

## Acknowledgments

We are grateful to Nathan Young and Rich Schneider for generously sharing their time and expertise related to geometric morphometrics, craniofacial biology, and avian embryology. We thank all past and present members of the Shapiro Lab, including Rebecca Bruders, Alexa Davis, Emily Maclary, Anna Vickrey, and Ryan Wauer for help with animal husbandry, technical assistance, and advice. We are grateful to Emily Maclary for comments on the manuscript. We thank members of the Utah Pigeon Club and National Pigeon Association for sample contributions. We acknowledge the University of Utah Preclinical Imaging Core Facility, especially Tyler Thompson, for micro-CT imaging; the Center for High Performance Computing at the University of Utah for computing resources; the University of Utah High-Throughput Genomics Shared Resource for RNA library preparation and sequencing; and the University of Minnesota Genomics Core for GBS library preparation and sequencing.

## Author Contributions

Conceptualization, E.F.B. and M.D.S.; Methodology, E.F.B. and M.D.S.; Software, C.H. and M.Y; Investigation, E.F.B. and H.F.V.; Resources, C.H. and M.Y; Formal Analysis, E.F.B.; Writing – Original Draft, E.F.B. and M.D.S.; Writing – Review & Editing, E.F.B., H.F.V., and M.D.S.; Funding Acquisition, E.F.B. and M.D.S.; Visualization, E.F.B.; Supervision, M.D.S.

---

**Supplemental Figure 1.**
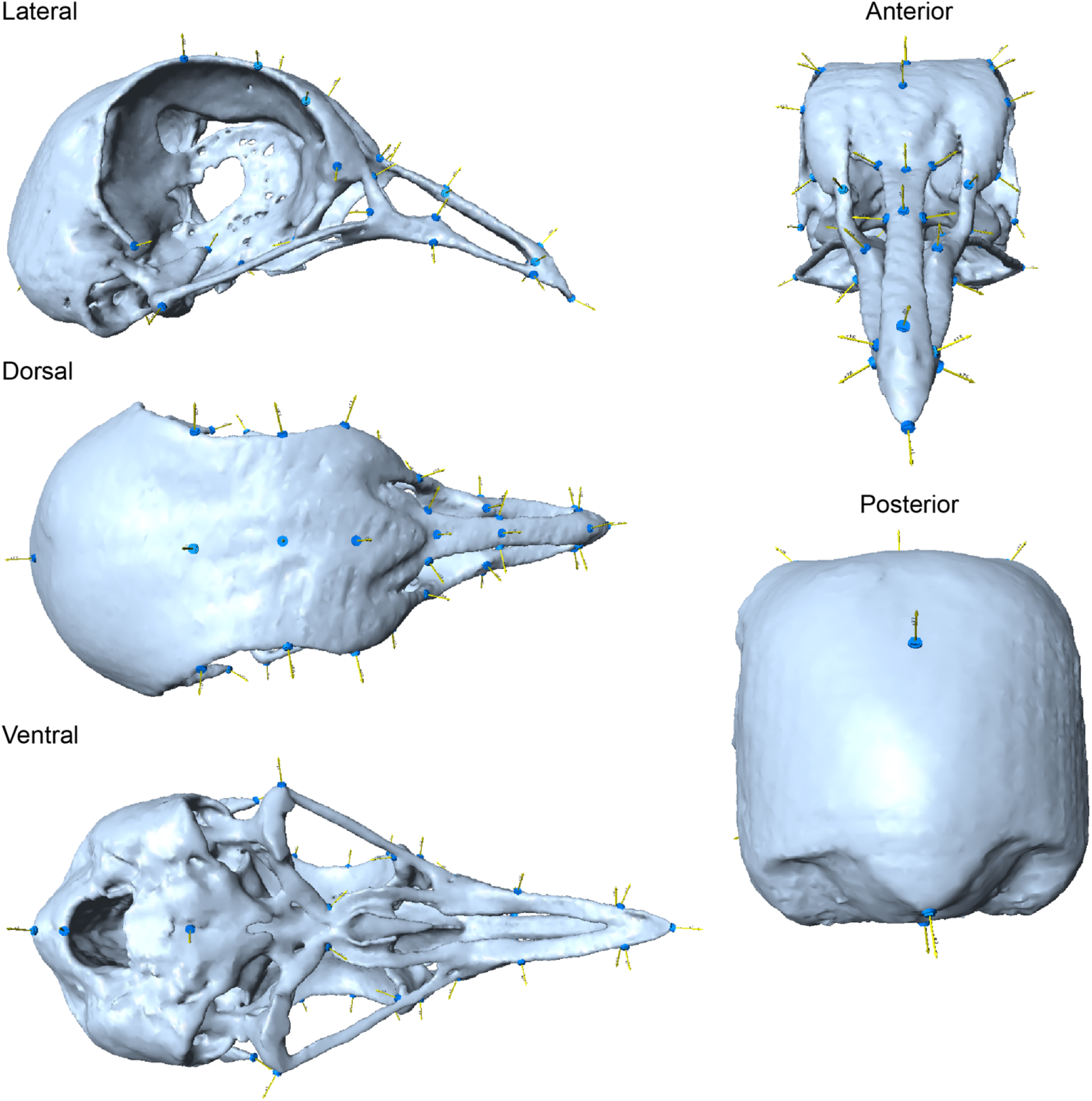
Pigeon craniofacial landmark atlas. Landmarks are indicated by blue discs.

**Supplemental Figure 2.**
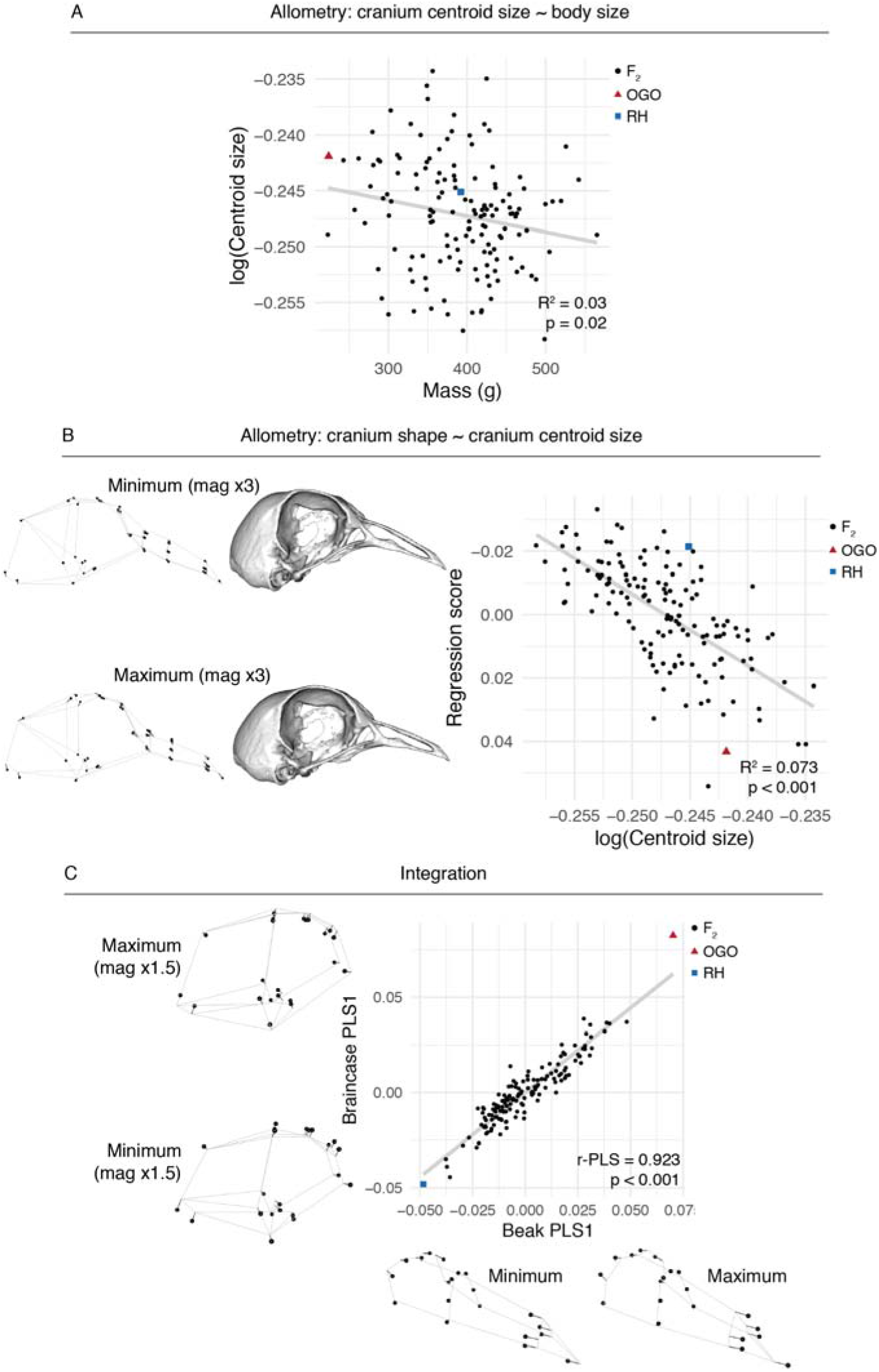
Allometry and integration in the RH x OGO cross. (A) Cranium centroid size ~ body size (mass) linear regression. (B) Cranium shape ~ centroid size linear regression. (B) Beak vs. braincase PLS1 shapes. Minimum and maximum shapes are depicted as wireframes and/or warped meshes along corresponding axis.

**Supplemental Figure 3.**
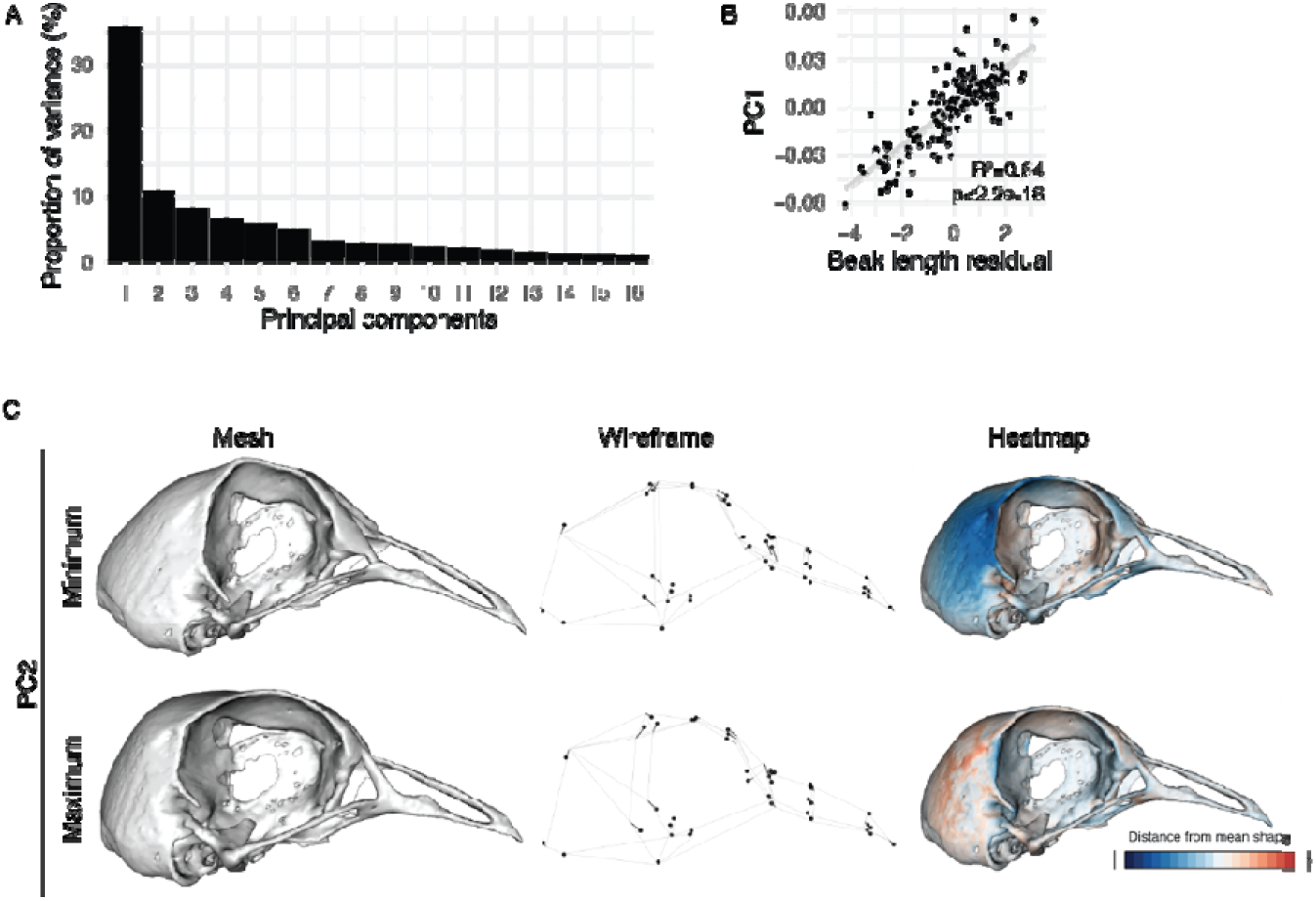
Geometric morphometric analysis of craniofacial skeleton shape in RH x OGO cross. (A) Principal components (PCs) that together account for 90% of shape variation in craniofacial skeleton. (B) Scatter plot of beak length residual vs. PC1 score for all RH x OGO F_2_ individuals. (C) Minimum and maximum PC2 shapes depicted as warped mesh (left), wireframe showing landmark displacement (center), or heatmap indicating regional shape changes (right). For mesh and wireframe models, shape change is magnified 1.5x to aid visualization.

**Supplemental Figure 4.**
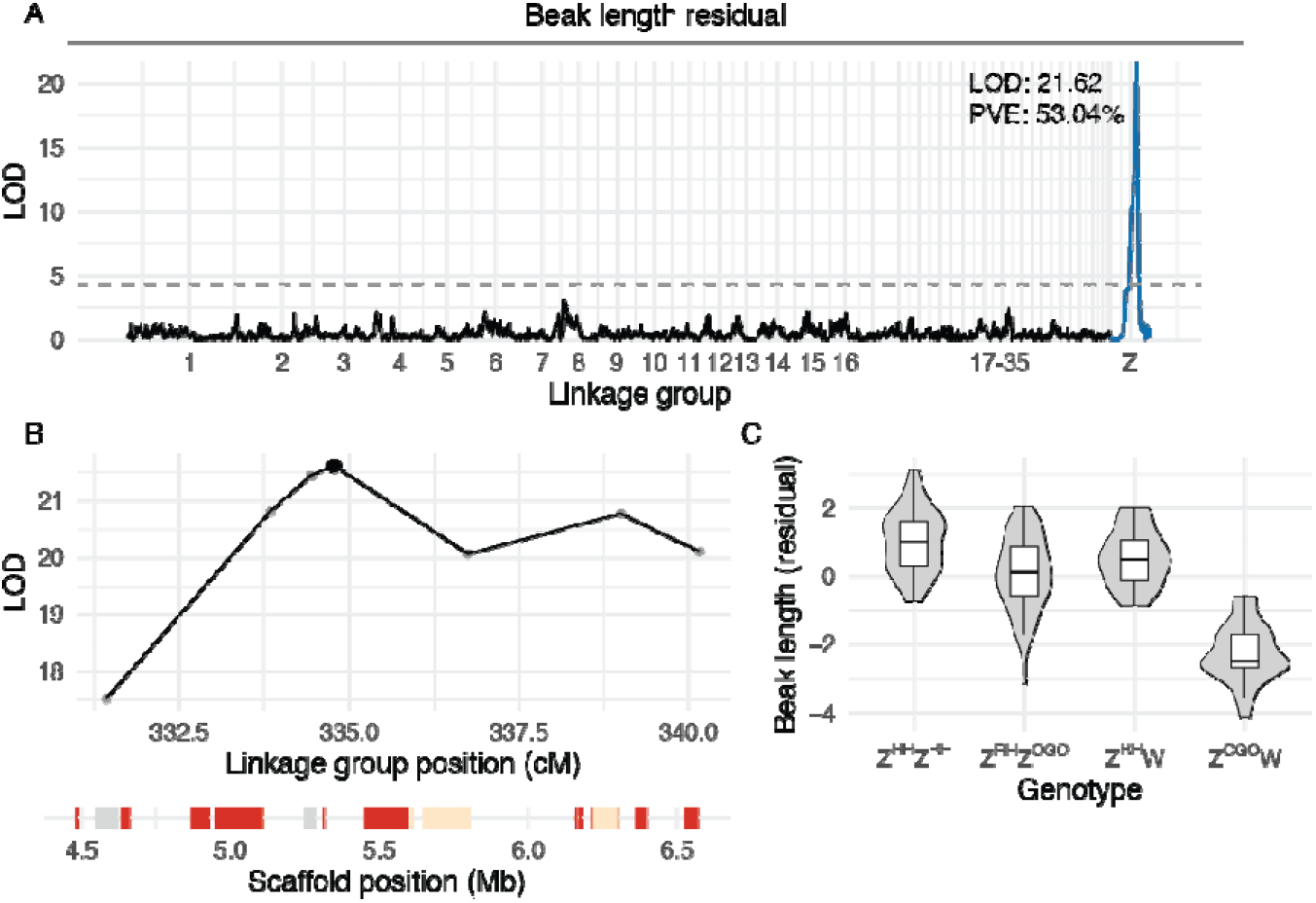
QTL scan using beak length residuals. (A) Genome-wide QTL scan for beak length using residuals from beak length ~ body mass linear regression. (B) LOD support interval is nearly identical to PC1 QTL interval (displayed in Figure 2D). Genes in interval on ScoHet5_445.1 are displayed at bottom and color-coded by expression level. (C) Plot of QTL effects, using peak marker highlighted by black dot in (B).

**Supplemental Figure 5.**
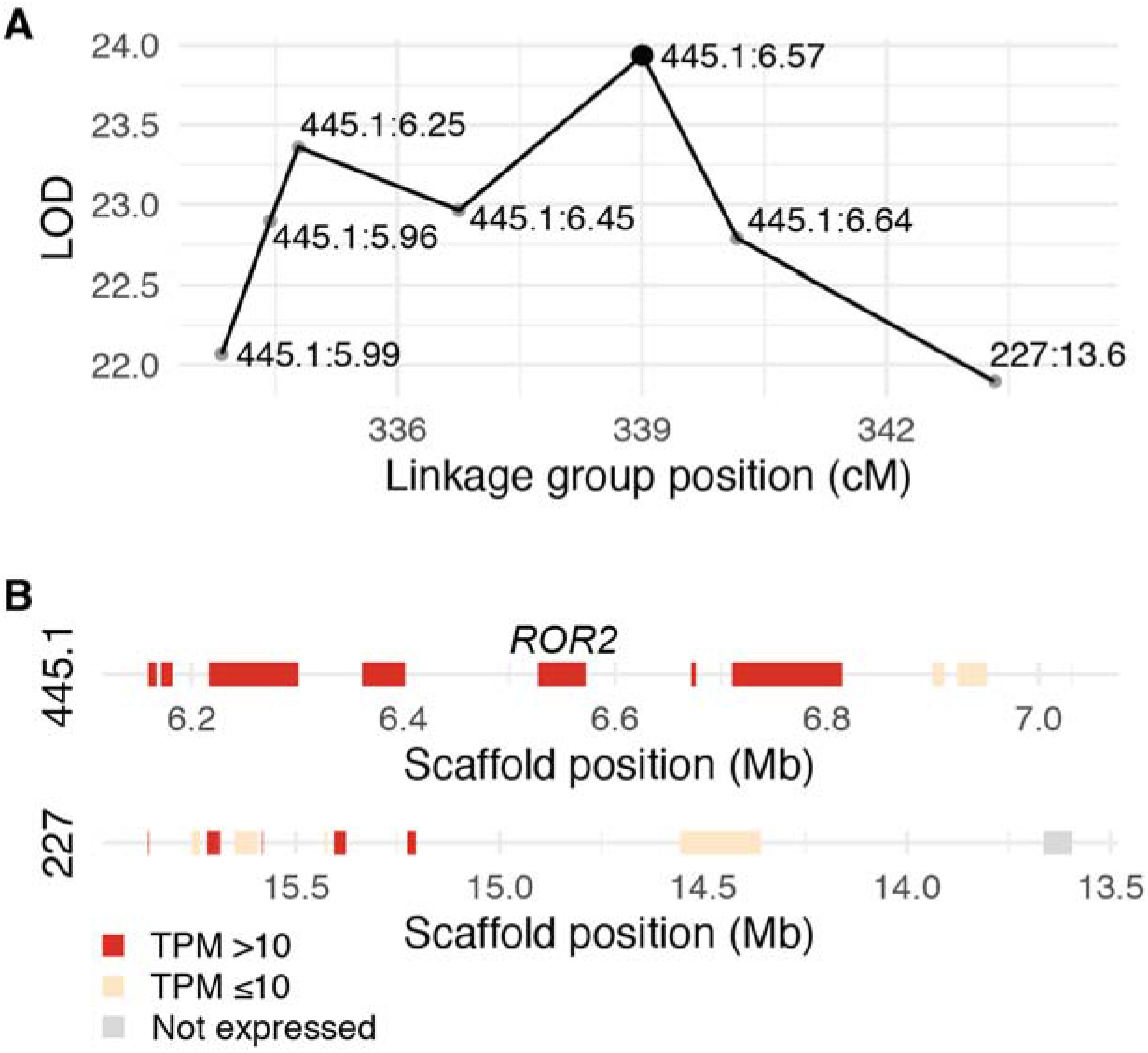
PC1 QTL support interval. (A) LOD support interval for QTL on Z. Markers in interval are denoted with dots and labeled with genomic scaffold name (ScoHet5_445.1 or ScoHet5_227) and position (in Mb); black dot indicates QTL peak marker (ScoHet5_445.1:6.57-Mb), which was used to estimate QTL effects. (B) Genes in QTL interval, color-coded by mRNA expression level in facial primordia derived from HH29 RH embryos. *ROR2* is located directly under QTL peak.

**Supplemental Figure 6.**
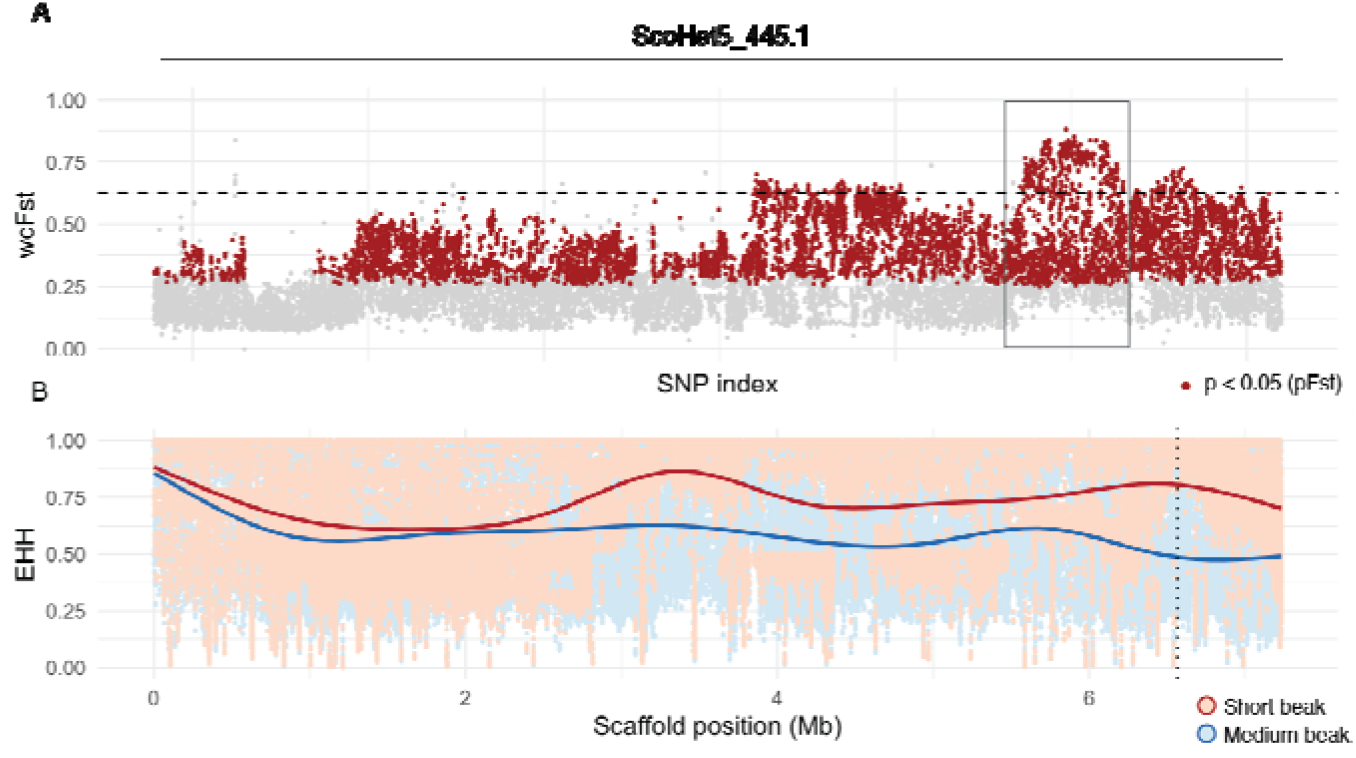
Allele frequency differentiation (F_ST_) and extended haplotype homozygosity (EHH) on scaffold ScoHet5_445.1 in short and medium/long beak pigeons. (A) F_ST_ on ScoHet5_445.1. Boxed region indicates ~293-kb peak region displayed in Figure 3D-E. (B) EHH on ScoHet5_445.1. Smoothed lines represent local regression fitting; dotted vertical line indicates position of ScoHet5_445.1:6568443.

**Supplemental Movie 1. PC1 shape variation.** Minimum to maximum, magnified 1.5x.

**Supplemental Movie 2. PC2 shape variation.** Minimum to maximum, magnified 1.5x.

**Supplemental Table 1. Description of skull and jaw landmarks.**

**Supplemental Table 2. Landmark pairs used for skull and jaw linear measurements.**

**Supplemental Table 3. Genes in the PC1 QTL interval.**

## STAR Methods

### Resource Availability

#### Lead Contact

Further information and requests for resources and reagents should be directed to and will be fulfilled by the Lead Contact, Michael Shapiro.

#### Materials Availability

Plasmids generated to synthesize RNA *in situ* hybridization probes against pigeon *ROR2* and *WNT5A* are available upon request.

#### Data and Code Availability

Whole genome sequencing and RNA-sequencing datasets generated for this study have been deposited to the NCBI SRA database under BioProject PRJNA680754. Additional short-beaked genomes are available under BioProject PRJNA513877 (SRR8420387-SRR8420391, SRR8420393, SRR8420394, SRR8420397).

### Experimental Model and Subject Details

#### Columba livia

Pigeons were utilized in accordance with protocols approved by the University of Utah Institutional Animal Care and Use Committee (protocols 10-05007, 13-04012, and 19-02011). Further information is provided throughout the Method Details section.

### Method Details

#### RH x OGO F_2_ intercross and 3D imaging

A F_2_ intercross was established between a male Racing Homer (RH) and a female Old German Owl (OGO). F_1_ hybrids (n=15) were interbred to generate a F_2_ mapping population. F_2_ offspring that reached 6 months of age (n=145) were euthanized and basic biometrics (e.g. mass) were recorded. All F_2_ offspring and cross founders were submitted to the University of Utah Preclinical Imaging Core Facility for micro-CT imaging. For each bird, a whole-body scan was performed on a Siemens Inveon micro-CT using the following parameters: voxel size=94 μ, photon voltage=80 kV, source current=500 μA, exposure time=200 ms. Scans were reconstructed using a Feldkamp algorithm with Sheep-Logan filter and a calibrated beam hardening correction.

#### Surface model generation and landmarking

Surface model generation and landmarking were performed as described ^24^. Briefly, a substack containing the cranium was extracted from the whole-body DICOM file stack in ImageJ v1.52q, exported as a NifTI file (*.nii), and imported into Amira v6.5.0 (Thermo Fisher Scientific). Using the Segmentation Editor threshold feature, the cranial skeleton was segmented from soft tissue and exported as a HxSurface binary (*.surf) file. Surface meshes were converted to Polygon (Stanford) ASCII (*.ply) files using i3D Converter v3.80 and imported into IDAV Landmark Editor v3.0 (UC Davis) for landmarking. We applied a set of landmarks (Supplemental Figure 1, Supplemental Tables 1-2) to the braincase (n=29 landmarks) and upper beak (n=20) of all F_2_ individuals and the cross founders. Landmark coordinates were exported as a NTsys landmark point dataset (*.dta) for geometric morphometric analysis.

#### Blood collection and genomic DNA extraction

Blood samples from adult pigeons used for whole-genome sequencing were collected at local pigeon shows, at breeder’s homes, or in the Shapiro lab loft. RH x OGO F_2_ offspring were bled at time of fledging (approximately 1 month of age). For each individual, a blood sample was collected from the brachial vein and stored in an EDTA-coated sample tube at −80°C. RNase-treated genomic DNA was extracted using a DNeasy Blood and Tissue Kit and eluted in Buffer EB (Qiagen).

#### Genotyping-by-sequencing (GBS) and linkage map assembly

Genomic DNA samples from RH x OGO cross founders and 171 F_2_ offspring were submitted to the University of Minnesota Genomics Core for GBS library prep and sequencing. Genomic DNA samples were digested with ApeKI (NEB), then ligated with T4 ligase (NEB) and phased adaptors with CWG overhangs. The ligated samples were purified with SPRI beads and amplified for 18 PCR cycles with 2X NEB Taq Master Mix to add barcodes. Libraries were purified, quantified, pooled, size selected for the 624-724 bp library region (480-580 DNA insert), and treated with ExoVII to remove any remaining single stranded material. The final pool was diluted to 1 nM for sequencing on the Illumina NovaSeq 6000 using single-end 1X100 reads. Target sequencing volume was ~4.75M reads/sample. Sequencing read quality was assessed with FastQC (Babraham Bioinformatics) and Illumina adapters were trimmed with Cutadapt ^55^. Reads were mapped to the Cliv_2.1 reference assembly ^56^ using Bowtie 2 ^57^. Genotypes were called using the Stacks v2.52 ref_map.pl program, which executes the Stacks pipeline programs gstacks and populations. The following options were passed to populations: -H -r 0.75 --map-type F_2_ --map-format rqtl.

The RH x OGO genetic map was constructed with the R package R/qtl v1.46-2 using genotype data from 171 F_2_ individuals. Because of differences in segregation patterns, autosomal and Z-linked scaffolds were assembled separately. For autosomal scaffolds, markers with identical genotypes or displaying segregation distortion (chi-square p < 0.005) were eliminated. Preliminary filtering was performed to remove markers missing in more than 20% (34/170) of F_2_ individuals. Pairwise recombination fractions were calculated and a preliminary genetic map was estimated using the *est.rf* and *est.map* functions, respectively. The *droponemarker* and *calc.errorlod* functions were used with the parameter (error.prob = 0.005) to identify problematic markers and likely genotyping errors, which were eliminated from the genetic map. Linkage groups were formed using the function *formLinkageGroups* with parameters (max.rf = 0.25, min.lod = 6). For the Z-chromosome, the same workflow was carried out, except that distorted markers were not removed. Preliminary marker ordering was done for all linkage groups using the *orderMarkers* function with the parameter (window size = 7). Final marker ordering was completed manually based on calculated recombination fractions and LOD scores. The *compareorder* function was used to test alternative marker orders; changes in marker ordering that resulted in an increased LOD score and decreased linkage group length were retained. The final RH x OGO genetic map is composed of 6128 markers (5553 autosomal, 575 Z-linked) on 35 linkage groups (34 autosomal, 1 Z-linked) with a genotyping rate of 90.1%.

#### Whole-genome resequencing

For the current study, we resequenced genomes for 33 pigeons from 24 short-beaked breeds: African Owl, Australian Tumbler, Berlin Short Face Tumbler, Budapest Tumbler, Canario Cropper, Chinese Nasal Tuft, Classic Old Frill, Damascene, Egyptian Swift, English Long Face Tumbler, English Short Face Tumbler, Granadino Pouter, Hamburg Sticken, Helmet, Italian Owl, Long Face Muff Tumbler, Nun, Old German Owl, Oriental Frill, Rafeno Pouter, Russian Tumbler, Taganrog Tumbler, Temeschburger Schecker, Uzbek Tumbler. We also resequenced 29 pigeons from 24 medium- or long-beaked breeds: Berlin Long Faced Tumbler, Dragoon, English Carrier, English Magpie, Scandaroon, Racing Homer, Danzig Highflier, Schalkaldener Mohrenkopf, Fairy Swallow, Hungarian Giant House Pigeon, Crested Saxon Field Color Pigeon, Saint, Franconian Trumpeter, American Highflier, Bokhara Trumpeter, Komorner Tumbler, Brunner Pouter, Mindian Fantail, Naked Neck, Turkish Tumbler, Norwich Cropper, Miniature American Crest, Rhine Ringbeater, Vienna MF Tumbler.

Genomic DNA samples were submitted to the High-Throughput Genomics and Bioinformatic Analysis Shared Resource at the University of Utah for library preparation and sequencing. DNA libraries were prepared using the Illumina TruSeq DNA PCR-Free Sample Preparation Kit with an average insert size of 350 bp. 125-cycle paired-end sequencing was performed on an Illumina HiSeq 2500 instrument (3-4 libraries/lane).

#### Embryonic tissue isolation and RNA extraction

Pigeon eggs were collected from Racing Homer (medium beak) and Oriental Frill (short beak) breeding pairs and incubated to embryonic day 6 (Hamburger-Hamilton (HH) stage 28-29, ^58^). Facial prominences that form the upper beak (frontonasal and maxillary, FNP+MXP) and lower beak (mandibular, MDP) were dissected and stored separately in RNAlater (ThermoFisher Scientific) at −80°C. Additional tissue was harvested from each embryo and used for DNA extraction and sex determination following a previously published PCR-based assay ^59^. Total RNA was extracted from embryonic tissue samples using the RNeasy Mini Kit with RNase-Free DNAse Set and a TissueLyser LT (Qiagen).

#### RNA-sequencing

Total RNA from FNP+MXP and MDP samples from HH28-29 female Racing Homer (n=5) and Oriental Frill (n=5) embryos was submitted to the High-Throughput Genomics and Bioinformatic Analysis Shared Resource at the University of Utah for library preparation and sequencing. RNA sample quality was assessed using the RNA ScreenTape Assay (Agilent). For each sample, a stranded sequencing library was prepared using the TruSeq Stranded mRNA Sample Prep Kit with oligo(dT) selection (Illumina). 125-cycle paired-end sequencing was performed on an Illumina HiSeq 2500 instrument (12 libraries/lane). An average of 23.4 million reads was generated for each sample.

#### ROR2 multiple sequence alignment

Amino acid sequences for vertebrate ROR2 and invertebrate ROR homologs were downloaded from Ensembl (ensembl.org) or NCBI (ncbi.nlm.nih.gov/gene). Clustal Omega multiple sequence alignments were performed and visualized with the R package msa v1.18.0 ^60^.

#### Whole-mount RNA in situ hybridization (ISH)

ISH probe templates were generated by PCR amplification of a portion of pigeon *ROR2* (692 bp amplicon) or *WNT5A* (783 bp amplicon) from a pooled cDNA library generated from HH21, HH25, and HH29 Racing Homer embryos using the following primer sets: ROR2-forward: 5’-GGAACCGACAGGTTCTACCA-3’, ROR2-reverse: 5’-TGCTTCGTCCATCTGAAGTG-3’, WNT5A-forward: 5’-CATAGTGGCTCTGGCCATTT-3’, WNT5A-reverse: 5’-CCCCGACTGTTGAGTTTCAT-3’. *ROR2* and *WNT5A* amplicons were cloned into pGEM-T Easy (Promega) and confirmed by Sanger sequencing. Antisense and sense RNA probes were generated by *in vitro* transcription as previously described ^61^. For *ROR2*, pGEM-*ROR2* was digested with NcoI or SalI and transcribed with SP6 or T7 RNA polymerase, respectively. For *WNT5A, pGEM-WNT5A* was digested with Kpn1 or Nco1 and transcribed with T7 or SP6 RNA polymerase, respectively.

Racing Homer embryos used for ISH were dissected from eggs at the desired embryonic stage and fixed overnight in 4% paraformaldehyde at 4°C on a shaking table. Embryos were subsequently dehydrated into 100% MeOH and stored at −20°C. Whole-mount ISH was performed following a protocol optimized for avian embryos (geisha.arizona.edu/geisha/protocols.jsp). For each experiment, antisense or sense probes were applied to stage-matched embryos.

### Quantification and Statistical Analysis

#### Linear measurement analysis

For each F_2_ individual and the cross founders, beak and braincase length were determined by calculating the linear distance between landmark pairs (beak: landmarks 1 and 2; braincase: landmarks 1 and 3, ^24^) using the *interlmkdist* function from the R package geomorph v3.3.1 ^62–64^. Raw beak length measurements were fit to a linear regression model (beak length ~ body mass) and residuals were calculated in R v3.6.3 ^65^.

#### Geometric morphometrics

Geometric morphometric analyses were performed in geomorph as described ^24^. The NTsys landmark point dataset was imported with the *readland.nts* function. Missing landmarks were estimated using the function *estimate.missing(method = “TPS”)*. Bilateral symmetry analysis was performed via the *bilat.symmetry(iter = 1)* function and the symmetrical component of shape variation was extracted. A Generalized Procrustes Analysis was performed using the *gpagen* function. To analyze allometry, a linear model (shape ~ centroid size) was fit using the *procD.lm* function and residuals were used for analysis of allometry-free shape. Principal components analysis was performed using the *gm.prcomp* function. Integration of beak and braincase shape was analyzed using the *two.b.pls* function.

Shape changes were visualized with geomorph and the R package Morpho v2.8 (https://github.com/zarquon42b/Morpho). The geomorph function *plotRefToTarget* was used to generate wireframes. Surface mesh deformations, heatmaps, and movies were generated in Morpho with the *tps3d, shade3d, meshDist*, and *warpmovie3d* functions. For all mesh-based visualizations, deformations were applied to a reference mesh, which was generated by warping a RH x OGO F_2_ mesh to the mean shape.

#### QTL mapping

QTL mapping was performed using the R package R/qtl v1.46-2 ^66^. Single-QTL genome scans were performed using the *scanone* function with Haley-Knott regression and sex as a covariate.

The 5% genome-wide significance threshold was calculated by running *scanone* with 1000 permutation replicates. For each QTL, the 1.5-LOD support interval was calculated with the *lodint* function, percent variance explained (PVE) was calculated with the *fitqtl* function, and QTL effects were estimated via the *plotPXG* function. We compared phenotypic means in RH x OGO F_2_ genotypic groups at peak markers via one-way ANOVA and Tukey Test for pairwise comparisons in R. Genes within QTL intervals were identified using a custom R script and visualized using the R packages ggplot2 v3.3.0 ^54^ and gggenes v0.4.0 (https://github.com/wilkox/gggenes).

#### Variant calling and comparative genomic analyses

Variant calling was performed with FastQForward ^67^, which wraps the BWA short read aligner ^68^ and Sentieon (sentieon.com) variant calling tools to generate aligned BAM files (fastq2bam) and variant calls in VCF format (bam2gvcf). Sentieon is a commercialized GATK equivalent pipeline that allows users to follow GATK best practices using the Sentieon version of each tool (broadinstitute.org/gatk/guide/best-practices and support.sentieon.com/manual/DNAseq_usage/dnaseq/). FastQForward manages distribution of the workload to these tools on a compute cluster to allow for faster data-processing than when calling these tools directly, resulting in runtimes as low as a few minutes per sample. Raw sequencing reads from 54 newly resequenced individuals (described in Whole-genome resequencing section) were aligned to the Cliv_2.1 reference assembly ^56^ using fastq2bam. Variant calling was performed for each newly resequenced individual, as well as 132 previously resequenced individuals ^27,30,50,69^, using bam2gvcf and individual genome variant call format (gVCF) files were created. Joint variant calling was performed on a total of 186 individuals using the Sentieon GVCFtyper algorithm. The resulting VCF file was used for all subsequent genomic analyses.

Genome-wide Weir and Cockerham’s F_ST_ (wcF_ST_) and probabilistic F_ST_ (pF_ST_) were calculated using the GPAT++ toolkit within the VCFLIB software library (github.com/vcflib) as previously described ^27,30,50,69^. Extended haplotype homozygosity (EHH) was calculated for genomic scaffold ScoHet5_445.1 using the GPAT++ sequenceDiversity tool. Putatively deleterious variants were identified using the Variant Annotation, Analysis, and Search Tool (VAAST2, ^36^), which was implemented as previously described ^27^. For all comparative genomic analyses, pigeons were binned into the following phenotypic groups:

Short beak (56 individuals, 31 breeds): African Owl, Australian Tumbler, Bacska Tumbler, Berlin Short Face Tumbler, Budapest Tumbler, Canario Cropper, Catalonian Tumbler, Chinese Nasal Tuft, Chinese Owl, Classic Old Frill, Damascene, Egyptian Swift, English Long Face Tumbler, English Short Face Tumbler, Granadino Pouter, Hamburg Sticken, Helmet, Italian Owl, Komorner Tumbler, Long Face Tumbler, Nun, Old German Owl, Oriental Frill, Portuguese Tumbler, Rafeno Pouter, Russian Tumbler, Spanish Barb, Syrian Dewlap, Taganrog Tumbler, Temeschburger Schecker, Uzbek Tumbler.

Medium or long beak (121 individuals, 58 breeds and feral): American Highflier, American Show Racer, Archangel, Armenian Tumbler, Berlin Long Face Tumbler, Birmingham Roller, Bokhara Trumpeter, Brunner Pouter, Carneau, Crested Saxon Field Color Pigeon, Cumulet, Danish Tumbler, Danzig Highflier, Dragoon, English Carrier, English Magpie, English Pouter, English Trumpeter, Fairy Swallow, Fantail, Feral, Franconian Trumpeter, Frillback, German Beauty, Hungarian Giant House Pigeon, Ice Pigeon, Indian Fantail, Iranian Tumbler, Jacobin, King, Lahore, Laugher, Lebanon, Marchenero Pouter, Mindian Fantail, Miniature American Crest, Modena, Mookee, Naked Neck, Norwich Cropper, Old Dutch Capuchin, Oriental Roller, Parlor Roller, Polish Lynx, Pomeranian Pouter, Pygmy Pouter, Racing Homer, Rhine Ringbeater, Runt, Saint, Saxon Monk, Saxon Pouter, Scandaroon, Schalkaldener Mohrenkopf, Shakhsharli, Starling, Turkish Tumbler, Vienna Medium Face Tumbler, West of England.

#### RNA-seq analysis

Analysis of RNA-seq data was performed as previously described ^70^. Briefly, sequencing read quality was assessed with FastQC (Babraham Bioinformatics). Illumina adapters were trimmed and reads were aligned to the pigeon Cliv_2.1 reference assembly (Holt et al., 2018) using STAR v2.5.0a ^71^ using the 2-pass mode. GTF annotation files were used to guide spliced read alignments. Mapped reads were assigned to genes using featureCounts from the Subread package version 1.5.1. Transcript abundance (TPM) was quantified using Salmon v1.3.0 ^72^. Differential expression analyses were performed with the R package DESeq2 version 1.12.4 (Love et al., 2014).

